# Basolateral amygdala and orbitofrontal cortex, but not dorsal hippocampus, are necessary for the control of reward-seeking by occasion setters

**DOI:** 10.1101/2022.07.28.501885

**Authors:** Kurt M. Fraser, Patricia H. Janak

## Abstract

Reward-seeking in the world is driven by cues that can have ambiguous predictive and motivational value. To produce adaptive, flexible reward-seeking it is necessary to exploit occasion setters, other distinct features in the environment, to resolve the ambiguity of Pavlovian reward-paired cues. Despite this, very little research has investigated the neurobiological underpinnings of occasion setting and as a result little is known about which brain regions are critical for occasion setting. To address this, we exploited a recently developed task that was amenable to neurobiological inquiry where a conditioned stimulus is only predictive of reward delivery if preceded in time by the non-overlapping presentation of a separate cue - an occasion setter. This task required male rats to maintain and link cue-triggered expectations across time to produce adaptive reward-seeking. We interrogated the contributions of the basolateral amygdala and orbitofrontal cortex to occasion setting as these regions are thought to be critical for the computation and exploitation of state value, respectively. Reversible inactivation of either structure prior to the occasion-setting task resulted in a profound inability of rats to use the occasion setter to guide reward seeking. In contrast, inactivation of the dorsal hippocampus, a region fundamental for context-specific responding was without effect nor did inactivation of the basolateral amygdala or orbitofrontal cortex in a standard Pavlovian conditioning preparation affect conditioned responding. We conclude that neural activity within the orbitofrontal cortex and basolateral amygdala circuit is necessary to update and resolve ambiguity in the environment to promote cue-driven reward-seeking.

## INTRODUCTION

A given reward-paired cue may have myriad relationships with rewards, leading to difficulty ascertaining its motivational and predictive value at any given moment. To overcome this uncertainty, it is critical to make use of other environmental stimuli that resolve ambiguity about reward-predictive cues and promote flexible reward-seeking. These ambiguity-resolving cues are often referred to as occasion setters, historically termed ‘features’, as they act to set the occasion for reward-seeking but do not drive reward-seeking on their own nor acquire a direct relationship with reward (Holland, 1991, 1992; Fraser and Holland, 2019; Schmajuk and Holland, 1998; Trask et al., 2017). Occasion setters are thus a unique class of Pavlovian cues that can powerfully modulate the predictive and motivational value of a traditional conditioned stimulus, historically termed the ‘target’, without engendering many of the properties associated with conditioned stimuli. Despite this, little research has explored occasion setting and its relationship to reward-seeking in comparison to standard Pavlovian conditioning preparations where conditioned stimuli have absolute relations with the presence or absence of reward. As a result, very little is known about the neurobiological basis of occasion setting despite its implications across a wide spectrum of neuropsychiatric disorders like schizophrenia and addiction (Lubow and Gewirtz, 1995; Anagnostaras and Robinson, 1996; Anagnostaras et al., 2002; Ramos et al., 2002; Valyear et al., 2017).

The basolateral amygdala (BLA) and orbitofrontal cortex (OFC) are reciprocally connected structures that are critical for exploiting acquired knowledge about reward-predictive cues to update and guide reward-seeking (Price, 2007; Sharpe and Schoenbaum, 2016). Damage to either structure does not produce a drastic impairment in simple Pavlovian conditioning, but impairment is revealed in a variety of tasks in which acquired cue-based information must be exploited to produce adaptive behavior (Hatfield et al., 1996; Schoenbaum et al., 1998; Burns et al., 1999; Gallagher et al., 1999; Parkinson et al., 2000; Holland et al., 2001, 2002; Setlow et al., 2002; Pickens et al., 2003, 2005; Holland and Gallagher, 2003; Schiller and Weiner, 2004; McDannald et al., 2005, 2014; Corbit and Balleine, 2005; Ostlund and Balleine, 2007; Izquierdo and Murray, 2007; Ambroggi et al., 2008; Ishikawa et al., 2008; Johnson et al., 2009; Jones et al., 2012; Moorman and Aston-Jones, 2014; Lopatina et al., 2015; Lichtenberg et al., 2017; Stolyarova and Izquierdo, 2017). In addition, disruptions in the BLA impair cue-related neural activity in the OFC and damage within the OFC impairs cue-related neural activity within the BLA suggesting communication between these regions is necessary for the proper encoding of cue-reward relationships (Schoenbaum et al., 2003; Saddoris et al., 2005; Lucantonio et al., 2015; Saez et al., 2015, 2017). Within this circuit, the BLA has been proposed to encode the current state value of the environment which is then conveyed to the OFC to integrate into a broader state space that represents multiple features relevant for the current task, that can be used to relay information to downstream targets to guide behavior (Belova et al., 2008; Morrison and Salzman, 2010; Parkes and Balleine, 2013; Wilson et al., 2014; Stalnaker et al., 2015; Sharpe and Schoenbaum, 2016; Wikenheiser and Schoenbaum, 2016; Lichtenberg et al., 2017). We hypothesized that activity within either structure would be critical for occasion setting, as this task includes cue-driven state transitions that require the constant maintenance and updating of state value to guide reward-seeking (Fraser and Janak, 2019). To test this, we trained rats in an occasion setting task and then reversibly inactivated either the BLA or the OFC. These effects were contrasted with inactivation of the dorsal hippocampus (DH), a region implicated in the processing of spatial contexts and in the control of behavioral responding by physical settings (Fuchs et al., 2005; Allen et al., 2016). We reveal that neural activity in both BLA and OFC is necessary for linking cue-triggered expectations across time to resolve ambiguity about a conditioned stimulus and produce adaptive reward-seeking.

## MATERIALS AND METHODS

### Subjects

Subjects were 73 experimentally naïve male Long-Evans rats (Envigo, Frederick, MD) approximately 60 days of age and weighing 250-300 g on arrival. Upon arrival, rats were single-housed in ventilated cages with *ad libitum* access to food and water in a temperature-and humidity-controlled room and maintained on 12:12 light/dark cycle (lights on at 07:00). After one week of acclimation to the colony room, feeding was restricted to maintain weights at ∼95% of *ad libitum* feeding weights. Food restriction was maintained for the duration of the experiment except for the post-surgical recovery period. All behavioral procedures took place between 08:00 and 12:00. All procedures were approved the Animal Care and Use Committee at Johns Hopkins University and followed the recommended guidelines in the Guide for the Care and Use of Laboratory Animals: Eighth Edition, revised in 2011.

### Apparatus

Behavioral testing occurred in ten identical Med Associates conditioning chambers (St. Albans, VT) housed in sound- and light-attenuating cabinets and controlled by a computer running MedPC IV software. In the center of one wall was a fluid receptacle that was located in a recessed port. On the opposite wall near the ceiling of the chamber was a white houselight (28 V) and to the right of the houselight was a white-noise generator (10-25 kHz, 20 dB). Outside of the behavioral chamber but within the sound- and light-attenuating cabinet was a red houselight (28 V) that provided background illumination during each behavioral session. Fluids were delivered to the port via tubing attached to a 60 mL syringe placed in a motorized pump located outside each cabinet.

### Pretraining

For at least two days prior to any procedures rats were extensively handled and acclimated to the experimenter. Rats were also given free access to 15% sucrose in water for 24 hours in their homecage to prevent any neophobia and accustom them to the future reward solution. At least one day after sucrose pre-exposure, pretraining was conducted in an approximately one-hour session in which rats were able to drink sucrose from the recessed reward port. During this session there were 80 2-second activations of the syringe pump (∼0.07 mL per delivery) on a variable time 60 s schedule (30-90 s range). All rats consumed all sucrose delivered in the pretraining session.

### Occasion Setting Task

Training in the occasion setting task took place in 3 distinct phases with each session lasting ∼2 hours on average with an average inter-trial interval of 3.3 minutes. There was one session a day. In the first phase, all trials (25 for BLA group, 30 for OFC, DH, and NAc groups) were reinforced presentations of the following sequence of events: occasion setter houselight for 5 s, a 5 s empty period, and 5 s of the conditioned stimulus white noise. Reward consisted of 5 s activation of the syringe pump resulting in delivery of ∼0.18 mL of 15% sucrose in the reward port occurred immediately upon termination of the conditioned stimulus white noise. Phase one training lasted for 4 sessions. In phase two, 40% of trials were reinforced as before, but the other 60% of trials were conditioned stimulus alone trials consisting of presentation of the white noise for 5 s with no reward (25 total trials for BLA group, 30 total trials for OFC, DH, and NAc groups). Phase two training lasted for 6 sessions. In the final phase of training, rats were exposed to the full occasion setting task (Figure 1A) consisting of 10 reinforced trials, 10 conditioned stimulus alone trials, and 10 occasion setter alone trials where the houselight was activated for 5 s with no subsequent stimuli or reward. Note that this task is purely Pavlovian; there was never a response requirement. Training in the final phase lasted for 4 sessions prior to surgery.

**Figure 1.**
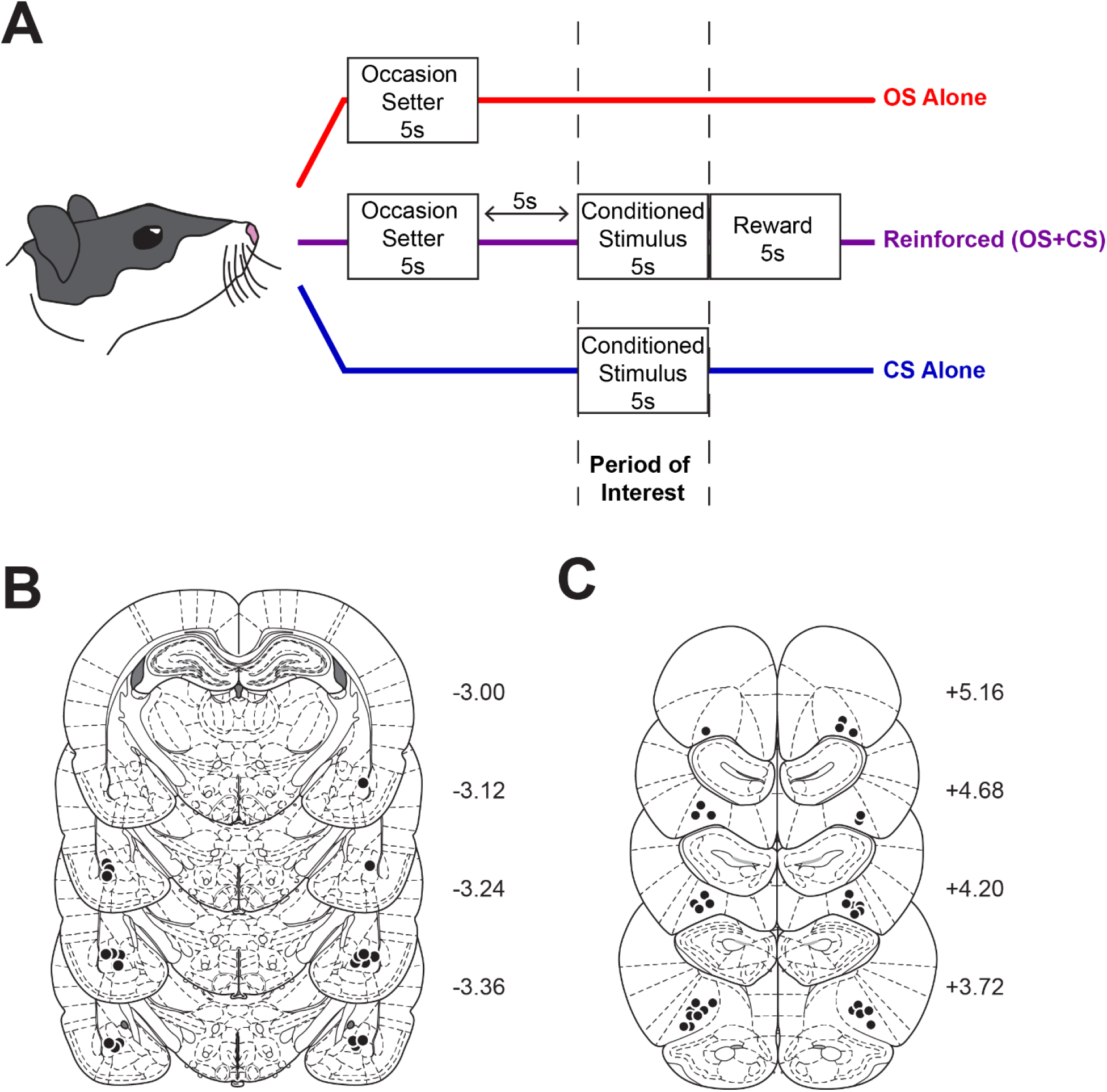
Experimental design and histological verification of microinjector tips. ***A***, Schematic of the occasion setting task. ***B***, Cannulae placements for rats in the BLA group (n=10). ***C***, Cannulae placements for rats in the OFC group (n=16). Numbers indicate distance from bregma in millimeters and the coronal sections were obtained from (Paxinos and Watson, 2007).

### Simple Conditioning Task

To assess the impact of BLA or OFC inactivation on Pavlovian conditioned responding separate groups of rats were instead trained in a task where a 5 s white noise was followed by 5 s activation of the reward pump containing 15% sucrose. The number of trials (30 per session) and intertrial interval (average 3.3 min) matched that of the occasion setting task and rats were trained for 14 days prior to cannula implantation. Following recovery, rats were retrained for 4 days before microinfusion testing.

### Surgery

After 14 sessions of training in the occasion setting task rats were anesthetized with isoflurane (5% induction, 1-2.5% maintenance) and standard stereotaxic procedures were used to implant 22 gauge cannula (Plastics One, Roanoke, VA) 1 mm above the intended infusion site in the BLA (n=30; AP: -3.0 ML: ±5.1 DV: -7.4; with injector DV: -8.4), OFC (n=31; AP: +3.5 ML: ±2.6 DV: -4.5; with injector DV: -5.5), DH (n=12; AP: -3.7 ML: ±2.5 DV: -2.5; with injector DV: -3.5). After surgery, rats received injections of cefazolin (70 mg/kg, subcutaneous) to prevent infection and carprofen (5 mg/kg, subcutaneous) to relieve pain. At all times except for during infusions, dummy stylets were placed in each guide cannula. These coordinates were adapted from previous studies (Chaudhri et al., 2013; Keiflin et al., 2013; Lichtenberg et al., 2017). Rats were allowed to recover for one week after surgery during which they had *ad libitum* access to food and water and then were returned to food restriction. Following this period, rats were retrained in the final phase of the occasion setting or simple conditioning task for at least 4 sessions.

### Infusions and Test

Rats were accustomed to the handling and infusion procedure for 1-2 days prior to infusion by being transported to the procedure room after a training session, dummy stylets removed, a flat cut 28 gauge injector inserted into each cannula, and dummy stylets replaced. On the day prior to testing, a regular injector extending 1 mm past the guide cannula was used to confirm cannula patency. At test rats received infusions of either saline or a mixture of the GABA-B and GABA-A agonists, baclofen and muscimol (1.0 mM and 0.1 mM, respectively) infused in a volume of 0.3 µL over 1 minute. After 1 additional minute to allow for diffusion away from the infusion site, injectors were removed, dummy stylets were replaced, and rats returned to their homecage for 5-10 minutes before test. There were two tests for each rat, one in each condition, with at least one day of retraining without manipulation between. Test sessions were reinforced.

### Histology

At the end of behavioral procedures, locations of cannulae were confirmed using standard histological procedures. Brains were extracted and post-fixed in 4% paraformaldehyde in 0.1 M NaPB for at least 24 hours and then cryoprotected in 25% sucrose in 0.1 M NaPB; 50 µm coronal sections were mounted onto Fisher SuperFrost Plus slides, and stained with cresyl violet (FD Neurotechnologies; Ellicott City, MD). Microinjection sites were verified by mapping their locations onto images from a rat brain atlas (Paxinos and Watson, 2007). Nine rats were excluded from the BLA cannula group and three rats were excluded from the OFC cannula group as a result of misplaced cannulae.

### Experimental Design and Statistical Analysis

All data were visualized and analyzed in GraphPad Prism 7. For all hypothesis tests, α=0.05. The primary behavioral data of interest were the effects of treatment (saline vs baclofen/muscimol) on time in port during the conditioned stimulus period on each trial type (reinforced, conditioned stimulus alone, occasion setter alone) in the occasion setting task or merely time in port between conditions for the simple conditioning task. It is important to note that on occasion setter alone trials, there is no conditioned stimulus presented, but the period analyzed is the corresponding 5 s period when the conditioned stimulus was presented on reinforced trials. These data were analyzed using a repeated measures ANOVA with within-subject factors of treatment and trial type. Difference scores were calculated to quantify each subject’s individual discrimination performance between time in port during the conditioned stimulus period on reinforced and either conditioned stimulus alone or occasion setter alone trials. These data were analyzed with a repeated measures ANOVA with treatment and discrimination (conditioned stimulus alone vs occasion setter alone) as within-subject factors. The observed effects were similar regardless of if time in port was normalized to the 10 s period prior to any stimulus onset. As a result, we present percent time in port data without normalization since there are overall increases in port time throughout the session that could increase the likelihood of observing decreases in responding with normalized measures. Intertial time in port and port entries were analyzed with two-tailed paired t-tests. When appropriate, post hoc comparisons were made with Tukey’s or Bonferroni’s procedure. Behavioral data for the BLA and OFC experimental groups was collected in separate sequential experiments and therefore were analyzed separately.

## RESULTS

To assess the contributions of the BLA and OFC to occasion setting male rats were trained in an occasion setting task and implanted with cannula over either the BLA or OFC. In this task, rats received deliveries of sucrose if the occasion setter (OS) and conditioned stimulus (CS) were linked in time with a 5 s gap between their presentations, whereas presentations of the OS alone or CS alone were non-reinforced (Figure 1A). As a result of this configuration of cues, rats exhibit higher conditioned approach behavior to the reward port on reinforced trials during CS presentation, than on CS alone trials, or during the CS period on OS Alone trials even though the CS was not presented. Overall, this general pattern of reward-seeking is consistent with previous reports of occasion setting in freely moving rats (Meyer and Bucci, 2016a, 2017). To determine the role of BLA and OFC in this differential responding during the CS period, rats received reversible inactivation of either structure with a mixture of the GABA-A and GABA-B agonists, muscimol and baclofen, or saline vehicle infusions in a counterbalanced manner before a reinforced session identical to that of the final phase of the occasion setting task. Only rats with microinjector tips verified within the bilateral BLA (n=10) or OFC (n=16) were included for analyses and their locations are depicted in Figure 1B and Figure 1C.

### Reversible inactivation of the BLA impairs occasion setting

Under control conditions (saline infusion), rats exhibited discriminated reward-seeking, as evidenced by heightened time in the reward port on reinforced trials relative to CS alone and relative to OS alone trials, and inactivation of the BLA abolished this pattern such that rats failed to discriminate among the three trials types, responding equivalently during the CS period regardless of the prior presentation of the OS or not (Figure 2A). These observations are confirmed by a main effect of trial type (F_(2,18)_=9.231, p=0.0017) and treatment (F_(1,9)_=22.6, p=0.001), as well as an interaction of treatment and trial type (F_(2,18)_=15.42, p=0.0001). While responding during the CS was greater following the OS relative to both CS alone (p=0.041) and OS alone trials (p<0.0001), following BLA inactivation rats did not discriminate among trial types, resulting in a similar, low level of reward seeking between reinforced and CS alone trials (p=0.9995) and OS alone trials (p=0.9999). In addition, comparing responding within trial type, inactivation of the BLA significantly reduced reward-seeking during the CS period on reinforced (p<0.0001) and CS alone trials (p<0.0001) compared to saline infusions. Conditioned approach to the food cup was already low during OS alone trials, given there was no stimulus present during the CS period, and this was not significantly reduced by BLA inactivation (p=0.4583). These effects are also evident when analyzing individual rats’ discrimination scores that quantify the difference in responding to the CS on reinforced trials vs either the CS alone or OS alone trials in the occasion setting task, with higher numbers reflecting better discrimination. Inactivation decreased the mean discrimination scores in both cases (Figure 2B; main effect of treatment: F_(1,9)_=16.42, p=0.0029; post hoc CS alone discrimination: p=0.0246; post hoc OS alone discrimination: p=0.0040).These findings collectively indicate that inactivation reduced the ability to use the occasion setter to produce adaptive reward-seeking. We also examined total time in the reward port and total port entries to examine whether inactivation generally impaired activity during the 2 hour session. Inactivation did not significantly alter intertrial port entries (Figure 2C; t_(9)_=1.868, p=0.0946) or time in the reward port (Figure 2D; t_(9)_=1.783, p=0.1083) indicating that decreased behavior during the CS period cannot be attributable to activity impairment. Rather, the findings suggest that rats failed to organize their reward-seeking appropriately in response to the presented stimuli. In addition, all rats consumed all sucrose delivered during both test sessions, indicating that even the presence of reward was not able to overcome the ability of BLA inactivation to disrupt occasion setting.

**Figure 2.**
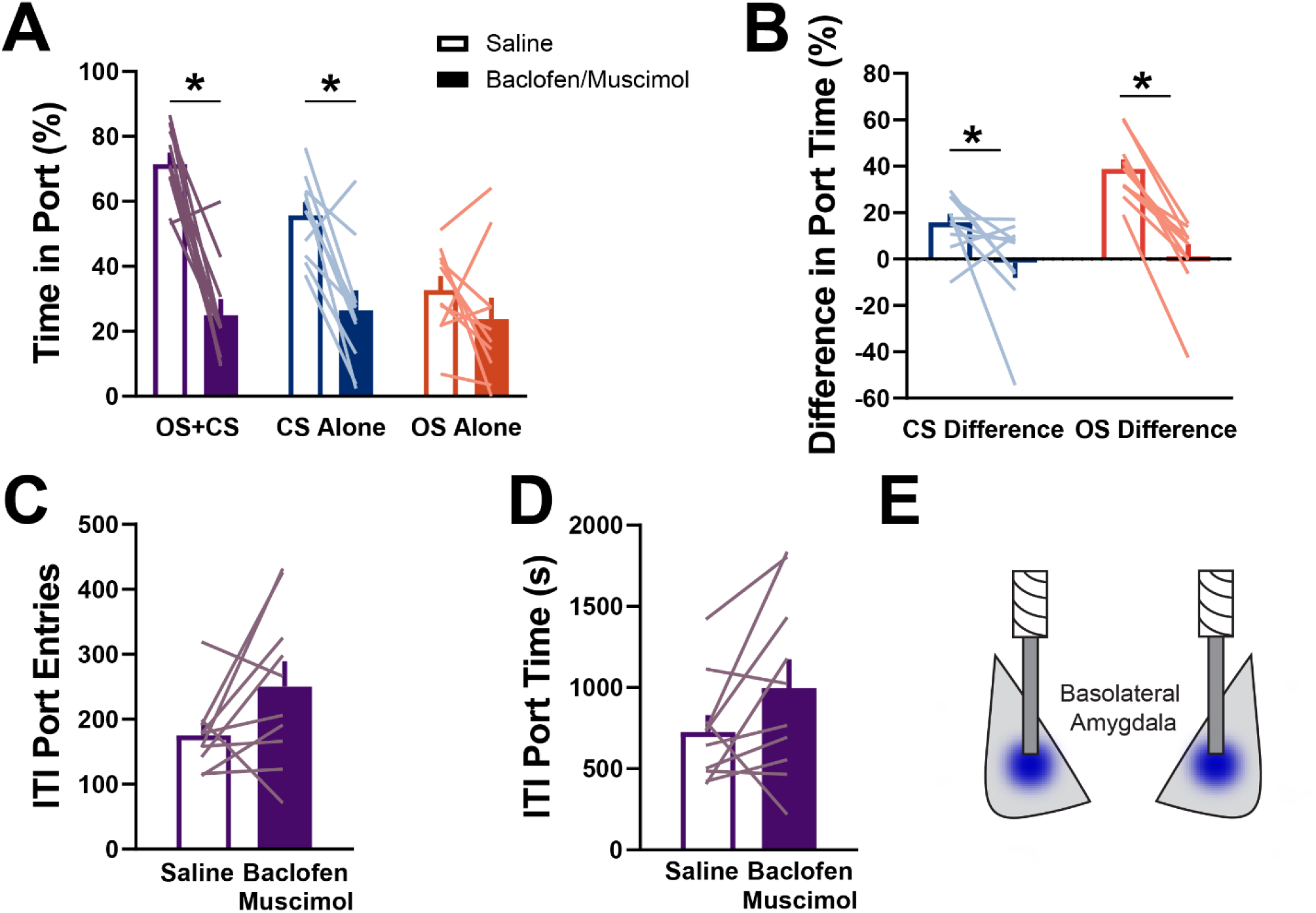
Reversible inactivation of the basolateral amygdala impairs occasion setting. ***A***, Average time in port during the conditioned stimulus period for both test sessions. ***B***, Individual differences in time in port during the conditioned stimulus period on reinforced trials minus either conditioned stimulus alone or occasion setter alone trials. ***C***, Intertrial port entries during the behavioral session. ***D***, Intertrial port time during the behavioral session. For all figures bars indicate mean ± SEM. Empty bars represent data from the saline session and filled bars from the inactivation session. Individual rats are overlaid and represented by the colored lines. OS, occasion setter. CS, conditioned stimulus. *p<0.05.

### Reversible inactivation of the OFC impairs occasion setting

In a separate group of rats, we tested whether inactivation of the OFC would alter occasion setting given its reciprocal connections with the BLA. Notably, we found that inactivation of the OFC produced a strikingly similar pattern of impairment in the occasion setting task (Figure 3A; main effect of trial type: F_(2,30)_=12.01, p=0.0001; main effect of treatment: F_(1,15)_=25.95, p=0.0001; interaction of treatment and trial type: F_(2,30)_=7.486, p=0.001). As above, rats exhibited discriminative reward-seeking in the task by using the occasion setter to increase reward-seeking on reinforced trials relative to CS alone (post hoc comparisons of reinforced versus CS alone: p<0.0001) and OS alone trials (p<0.0001) following saline infusions into the OFC. Similar to the findings above, inactivation of the OFC eliminated the ability of rats to use the occasion setter to guide reward-seeking and produced an overall low level of conditioned behavior during the test session (Figure 3A) as evidenced by no significant difference in time in port during the CS period between reinforced trials and either CS alone (p=0.9999) and OS alone trials (p=0.4234). For OFC, inactivation significantly reduced responding on reinforced (p<0.0001) and CS alone trials (p=0.0045), but did not reduce the already low level of responding on OS alone trials (p=.459). As a result, rats’ individual discrimination between reinforced trials and CS alone or OS alone trials was abolished following reversible inactivation of the OFC (Figure 3B; main effect of treatment: F_(1,15)_=0.0024; post hoc CS alone discrimination: p=0.0024; post hoc OS alone discrimination: p=0.0001). We analyzed total time in port and total port entries to assess whether the findings could be attributed to an overall lack of engagement during the test session. As with BLA inactivation, OFC inactivation did not impair overall activity during the session, but instead significantly increased intertrial port entries (Figure 3C; t_(15)_=3.266, p=0.0052) and time in the reward port (Figure 3D; t_(15)_=5.793, p<0.0001). In addition, all rats consumed all sucrose delivered during the test sessions. Together these results indicate that inactivation of the OFC impaired the ability of rats to use an occasion setter to resolve ambiguity about conditioned stimuli to guide reward-seeking.

**Figure 3.**
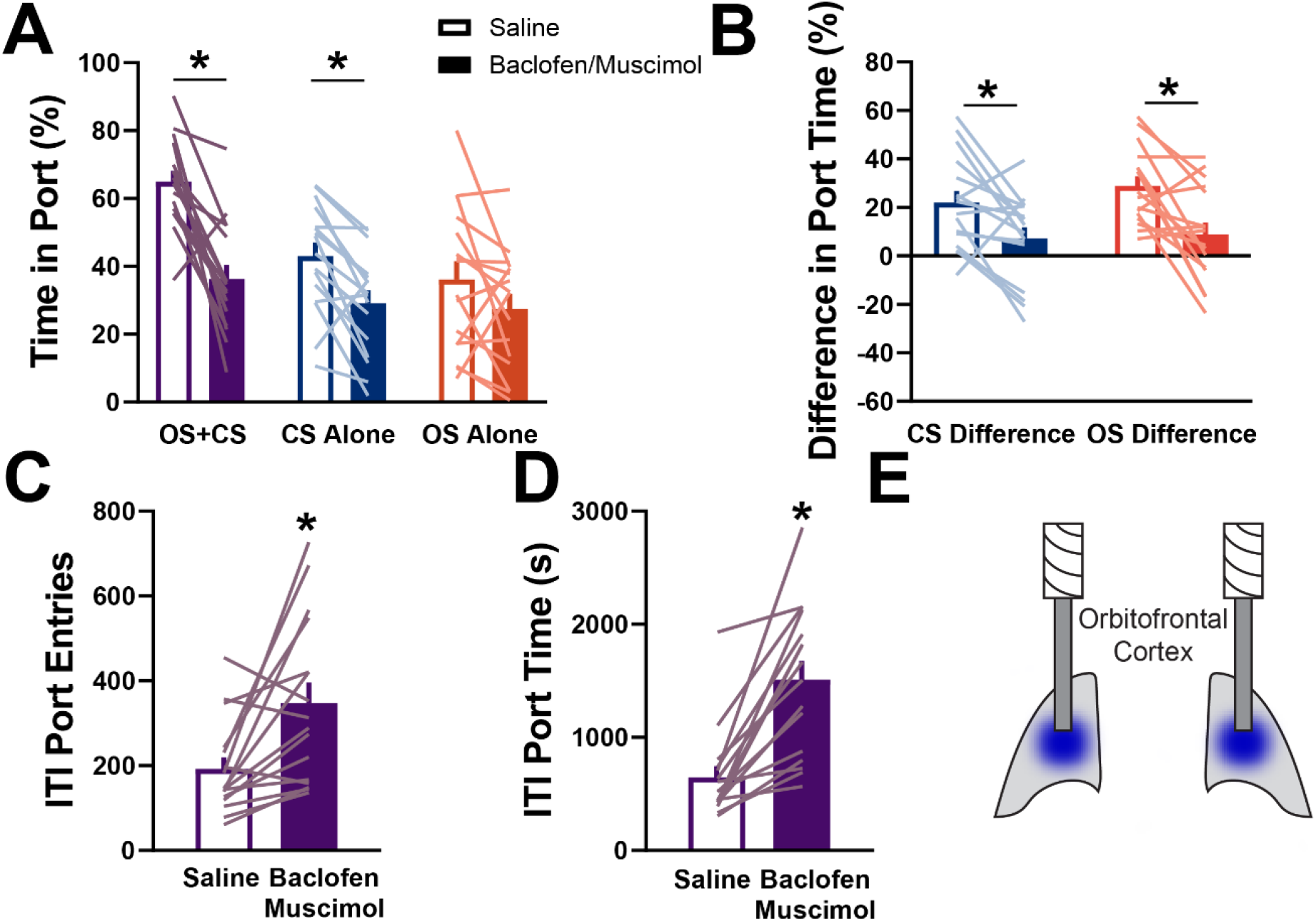
Reversible inactivation of the orbitofrontal cortex impairs occasion setting. ***A***, Average time in port during the conditioned stimulus period for each trial type. ***B***, Individual differences in time in port during the conditioned stimulus period on reinforced trials minus either conditioned stimulus alone or occasion setter alone trials. ***C***, Intertrial port entries during the behavioral session. ***D***, Intertrial port time during the behavioral session. For all figures bars indicate mean + SEM. Empty bars represent data from the saline session and filled bars data from the inactivation session. Individual rats are overlaid and represented by the colored lines. OS, occasion setter. CS, conditioned stimulus. *p<0.05.

### Reversible inactivation of the DH has no impact on occasion setting

We next asked whether the dorsal hippocampus, a region critical for the encoding of discrete locations in space, sequences of stimuli, and implicated in the control of responding by physical contexts may be critical in occasion setting. We trained 12 naïve rats in the occasion setting task as before and 2 rats were excluded for incorrect cannula placement (final n=10 for DH; placements in Figure 4E). In contrast to the impact of reversible inactivation with baclofen and muscimol in the BLA or OFC, there was no significant effect of inactivation of the DH on occasion setting assessed time in port during the conditioned stimulus across trials (Figure 4A; main effect of treatment F_(1,9)_=0.068, p=0.7997; main effect of trial type F_(2,18)_=16.03, p=0.0001; interaction of treatment and trial type F_(2,18)_=5.111, p=0.0175). Under both saline and inactivation rats exhibited more food cup responding on OS+CS trials relative to either CS Alone (saline p=0.0010; inactivation p<0.0001) and OS Alone trials (saline p<0.0001; inactivation p=0.0002), but there was no impact of inactivation on responding within any given trial (OS+CS p>0.9999; CS Alone p=0.6239; OS Alone p=0.2675). In agreement, analysis of the ability of each rat to discriminate between reinforced conditioned stimuli and non-reinforced trials revealed no significant effect of treatment (Figure 4B; F_(1,9)_=0.006, p=0.9386). Despite a lack of impact on occasion setting performance, inactivation of the DH significantly increased intertrial port entries (Figure 4C; t_(9)_=3.57, p=0.0060) but had no effect on the time rats spent in the port in the intertrial interval (Figure 4D; t_(9)_=1.536 p=0.1589). Together these indicate that the contributions of the DH to occasion setting are minimal, but that inactivation of the dorsal hippocampus can increase locomotor activity and disinhibit behavior in the absence of conditioned stimuli.

**Figure 4.**
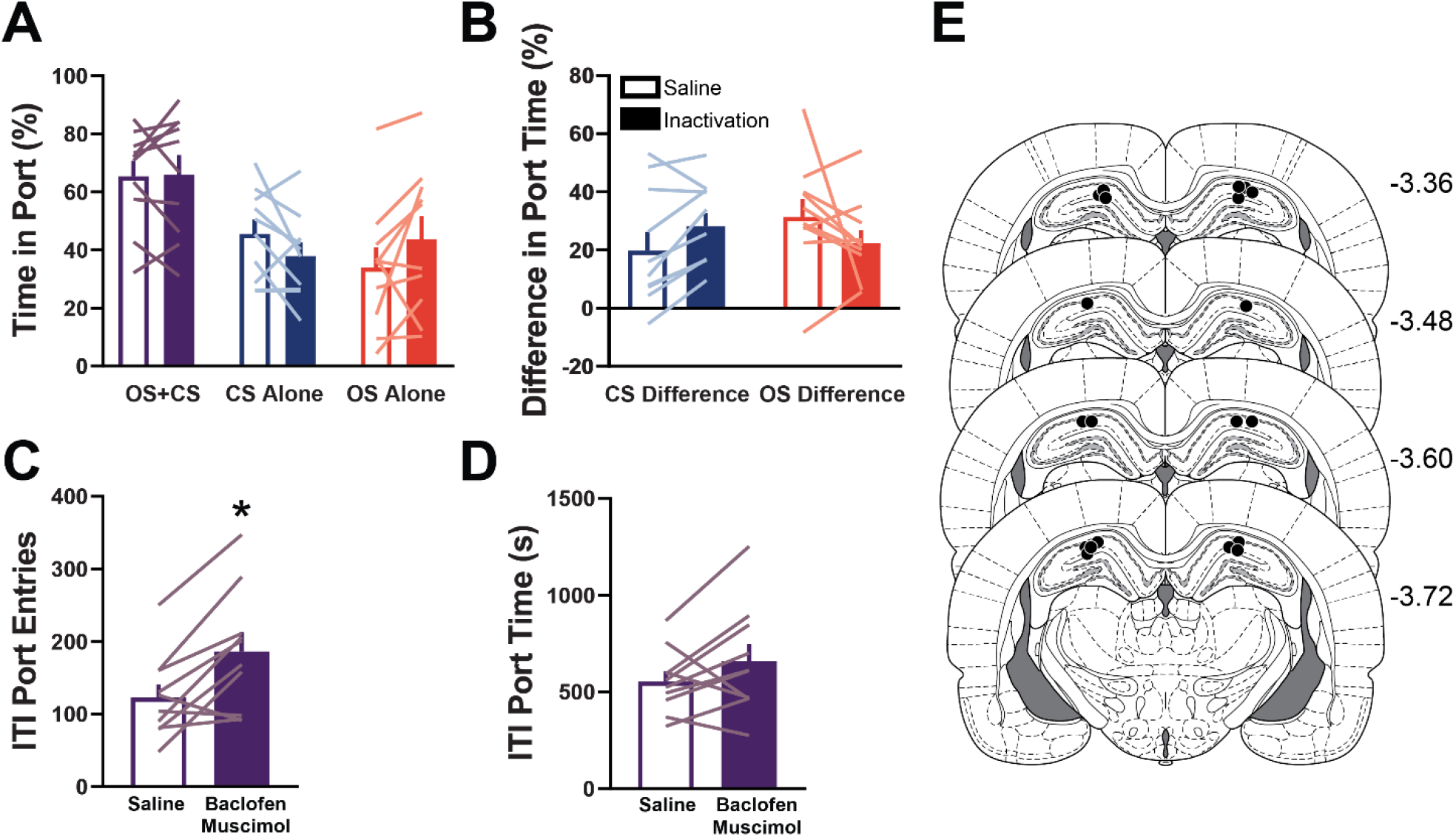
Dorsal hippocampus inactivation is without effect on occasion setting. ***A***, Average time in port during the conditioned stimulus period for each trial type. ***B***, Individual differences in time in port during the conditioned stimulus period on reinforced trials minus either conditioned stimulus alone or occasion setter alone trials. ***C***, Intertrial port entries during the behavioral session. ***D***, Intertrial port time during the behavioral session. ***E***, Cannulae placements for DH rats (n=10). Numbers indicate distance from bregma in millimeters and the coronal sections were obtained from (Paxinos and Watson, 2007). For all figures bars indicate mean + SEM. Empty bars represent data from the saline session and filled bars data from the inactivation session. Individual rats are overlaid and represented by the colored lines. OS, occasion setter. CS, conditioned stimulus. *p<0.05.

### Inactivation of either BLA or OFC fails to impair Pavlovian conditioned responding

Finally, we sought to confirm whether the impact of inactivation of the BLA or OFC on occasion setting could be explained by their requirement for conditioned responding. Despite evidence from multiple laboratories demonstrating that lesions of either structure or reversible inactivation are generally without effect on responding to Pavlovian cues, we wondered if the procedures employed here with a brief 5s cue followed by the non-overlapping delivery of reward might alter the involvement of either structure in Pavlovian conditioned approach. Rats were implanted with cannula over either the BLA (n=8 after exclusions) or the OFC (n=10 after exclusions) and trained for an identical number of sessions in a simple Pavlovian conditioning task where a 5s white noise was always followed by sucrose reward, with matched trial timing and trial number as the occasion setting task (Figure 5A). Reversible inactivation of the BLA after training for 18 sessions in this task was without effect on conditioned responding to the white noise (Figure 5B; t_(8)_=1.037, p=0.3344) but did significantly increase intertrial port time (Figure 5C; t_(8)_=2.476, p=0.0425). For OFC, there was similar a lack of impairment of reversible inactivation on conditioned responding to the white noise (Figure 5E; t_(11)_=2.012, p=0.0719), but inactivation did also significantly increase intertrial port time (Figure 5F; t_(11)_=2.868, p=0.0167). Ultimately, there is minimal contribution of either BLA or OFC to Pavlovian conditioned approach which suggests the extreme impairment in the occasion setting task is not due to their involvement in the generation of conditioned food cup approach.

**Figure 5.**
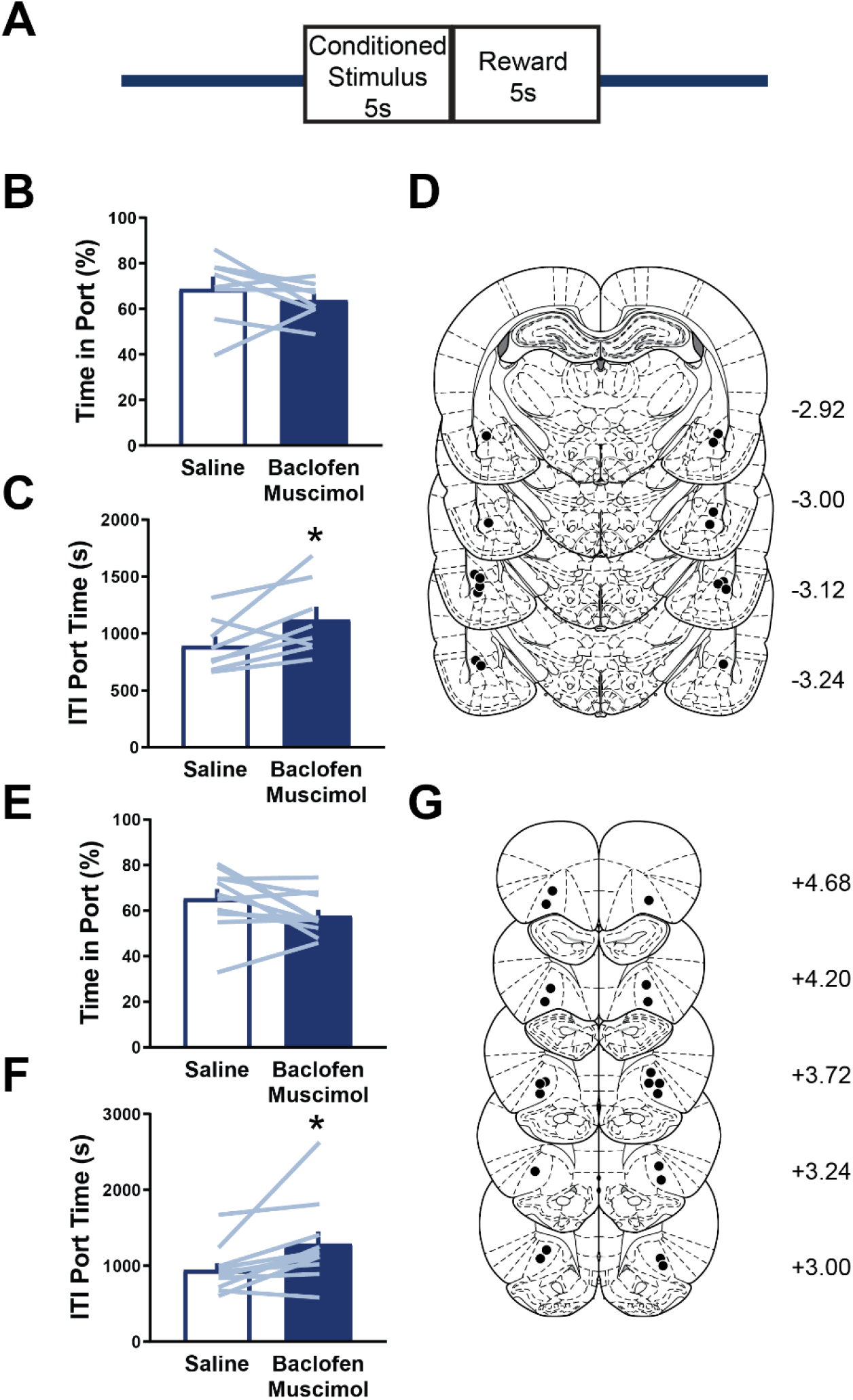
Inactivation of neither basolateral amygdala nor orbitofrontal cortex alters Pavlovian conditioned approach. ***A***, Schematic of task design where 30 trials per session were presented with each trial being the presentation of white noise for 5 s followed by 5 s activation of a pump containing 15% sucrose. ***B***, Time in port expressed as a percentage during the white noise. ***C***, Intertrial time in the port during the behavioral session. ***D***, Cannulae placements for BLA rats (n=8). ***E***, Time in port expressed as a percentage during the white noise. ***F***, Intertrial time in the port during the behavioral session. ***G***, Cannulae placements for OFC rats (n=11). Numbers indicate distance from bregma in millimeters and the coronal sections were obtained from (Paxinos and Watson, 2007). For all figures bars indicate mean + SEM. Empty bars represent data from the saline session and filled bars data from the inactivation session. Individual rats are overlaid and represented by the colored lines. *p<0.05.

## DISCUSSION

Resolving ambiguity about reward-predictive cues is an essential tool for survival, including the proper organization of reward-seeking behavior. However, in many experiments Pavlovian reward-predictive cues are deterministic with absolute relations with the presence or absence of reward and as a result are unambiguous. To better understand situations in which conditioned stimuli are ambiguous, we took advantage of an occasion setting task that required rats to use a discrete occasion-setting cue which signaled that if a conditioned stimulus was presented shortly after the occasion setter it would be followed by reward (Holland, 1992; Fraser & Holland 2019; Fraser & Janak 2019), while conditioned stimulus presentations not preceded by the occasion setting cue would not be followed by reward. We assessed the contributions of the BLA and OFC in regulating behavior under these conditions as these structures have been shown to be important for the generation and exploitation of cue-elicited expectations. We demonstrate that reversible inactivation of either structure produced a profound inability of rats to use an occasion setting cue to guide adaptive reward-seeking. Inactivation also increased overall time in the reward port, particularly following OFC inactivation, suggesting an inability in using cues to guide reward-seeking as opposed to a motor impairment. These data suggest activity within these reciprocally-connected structures, and likely communication within this circuit, is critical for resolving ambiguity about reward-predictive cues.

### Occasion setting mechanisms in conditioned reward-seeking

Despite extensive behavioral work, little is known with respect to the neural systems and circuits that are critical to occasion setting. Occasion setting is a rich and complex behavioral scenario that requires animals to maintain multiple cue-generated representations and link them in time to resolve the predictive and motivational meaning of a conditioned stimulus (Fraser and Holland, 2019; Fraser and Janak, 2019). In recent years there has been a renewed focus on negative occasion setting, a task in which the occasion-setting cue signals a conditioned stimulus will not be reinforced, thereby inhibiting reward-seeking (Holland, 1992; Meyer and Bucci, 2016b). These experiments have suggested that the OFC and nucleus accumbens, as well as the dorsal hippocampus, are involved in the acquisition of this behavior (Holland et al., 1999; Meyer and Bucci, 2016a; Shobe et al., 2017). Negative occasion setting differs in a significant number of ways from the positive occasion setting task that we used here, in which the occasion setting cue instructs the animal that it may soon encounter a conditioned stimulus that will spur reward-seeking (Holland, 1992; Meyer and Bucci, 2016b; Trask et al., 2017). Negative occasion setting requires behavioral inhibition, and negative occasion setting cues may act as conditioned inhibitors (Meyer and Bucci, 2017; Trask et al., 2017). Interestingly, evidence suggests distinct neural systems may underlie negative and positive occasion setting (Holland et al., 1999). In support of this distinction, we observed no impact of reversible inactivation of the dorsal hippocampus on positive occasion setting. To our knowledge ours is one of the first reports of a neurobiological manipulation to selectively affect positive occasion setting, laying a groundwork for future investigations into the neural circuits regulating the acquisition and expression of positive versus negative occasion setting.

### Basolateral amygdala and orbitofrontal cortex are critical for occasion setting

In many situations the BLA is not necessarily critical for the acquisition and expression of simple Pavlovian conditioned responses, but is necessary for exploiting and updating previously acquired cue-based information (Hatfield et al., 1996; Setlow et al., 2002; Holland and Gallagher, 2003; Pickens et al., 2003; Johnson et al., 2009; Chang et al., 2012). In the present study, in the absence of a functioning BLA, rats were not able to retrieve the appropriate motivational information associated with either cue in the occasion setting task (Averbeck and Costa, 2017), and as a result exhibited a flat, low-level of reward-seeking regardless of the cue presented. In contrast to studies of the BLA in context-based renewal of Pavlovian conditioned responding (Chaudhri et al., 2013; Millan et al., 2015; Sciascia et al., 2015), a situation in which a context must be used to inform reward-seeking, we observed significant impairments despite testing under reinforced conditions. This suggests perhaps that the nature of the occasion setting task, characterized by the need to rapidly update cue values every few minutes, more strongly engages the BLA. That inactivation of the BLA impairs occasion setting is in agreement with suggestions the BLA encodes state value, as occasion setting requires subjects to appropriately recognize and transition between cue-driven states (Morrison and Salzman, 2010). Neurons within the BLA respond to Pavlovian cues and to discriminative stimuli that signal the availability of an action to produce reward (Paton et al., 2006; Tye and Janak, 2007; Ambroggi et al., 2008; Morrison and Salzman, 2009; Shabel and Janak, 2009; Sangha et al., 2013), but whether or how they encode occasion setters themselves remains to be observed.

The OFC, like the BLA, is largely not necessary for simple Pavlovian conditioning, but is necessary for exploiting and updating acquired cue-based representations in new situations (Gallagher et al., 1999; Pickens et al., 2003, 2005; McDannald et al., 2005; Ostlund and Balleine, 2007; Chang, 2014). We found that inactivation of the OFC impaired occasion setting in well-trained subjects, suggesting an essential role of the OFC in the ongoing expression of this behavior. While neurons in the OFC acquire responses to reward-predictive stimuli, this information is encoded and used in a manner distinct from the BLA (Schoenbaum et al., 1998; Morrison and Salzman, 2011; Morrison et al., 2011; Takahashi et al., 2013; Moorman and Aston-Jones, 2014; Lopatina et al., 2017; Shobe et al., 2017). Neurons in the OFC have been argued to encode task space and ultimately construct a composite cognitive map of all possible states and their transitions (Wilson et al., 2014; Stalnaker et al., 2015; Wikenheiser and Schoenbaum, 2016). Accordingly, we hypothesize that inactivation of the OFC eliminated the ability of rats to use state value information, perhaps arising from the BLA (Schoenbaum et al., 2003; Sharpe and Schoenbaum, 2016; Lichtenberg et al., 2017), to maintain and/or transition in this state space appropriately, seemingly rendering all cues to have similar low significance to the subject as evidenced by the similar low level of responding across trials following OFC inactivation.

### BLA and OFC interactions in occasion setting

With the current approach we are unable to ascertain how communication between the OFC and BLA contributes to occasion setting. However, it is striking that the impairments in occasion setting observed after inactivation of either structure were almost identical. This suggests communication between OFC and BLA is critical for linking cue-triggered expectations across time to produce adaptive and flexible reward-seeking. Indeed, in an odor-guided decision-making task, neural encoding of cue-based information in the OFC is dependent on neural activity within the BLA (Schoenbaum et al., 2003), and encoding in the BLA is dependent on the OFC (Saddoris et al., 2005). A recent report suggests that the BLA encodes and retrieves cue-triggered expectations and the OFC exploits this information to guide decision-making (Lichtenberg et al., 2017). It is possible, however, that the identical effects we observed on occasion setting may be mediated by a shared downstream target of the BLA and OFC, such as the nucleus accumbens, but this remains to be explored (Heilbronner et al., 2016). Paired recordings in the OFC and BLA during this occasion setting task would give insight into the unique contributions of each structure and clarify the computations occurring in each, while future investigations using chemogenetic or optogenetic tools to restrict manipulations to BLA terminals in the OFC or OFC terminals in the BLA to resolve contributions of directionality in this circuit for occasion setting.

## Conclusions

The ability to resolve ambiguity surrounding reward-paired cues is essential for survival but often neglected in studies of Pavlovian conditioning. Occasion setting allows for a complex understanding of the dynamic regulation of cue-triggered reward-seeking by discrete events. The present data demonstrate that the BLA and OFC are essential neural substrates for exploiting occasion setting cues to produce flexible conditioned reward-seeking. Excessive and inappropriate pursuit of rewards is a hallmark of neuropsychiatric disorders, like addiction, that may arise from deficits in occasion setting processes, and a better understanding of the neural and behavioral mechanisms of occasion setting could provide insights into new clinical interventions.

## Acknowledgements

This work was supported by NIH Grants DA035943 to PHJ and DA046136 to KMF. We thank Alex Haimbaugh and Erin Kong for excellent technical assistance. We would also like to thank Dr. Peter Holland and members of the Janak Lab for their comments, input, and support. We would like to acknowledge the immense impact Dr. Nadia Chaudhri had as an inspiration for this line of research.

## Notes

**Conflict of Interest:** The authors declare no competing financial interests.

### Competing Interest Statement

The authors have declared no competing interest.

